# A microbiome-derived nutrient underlies tyrosine metabolism and predator avoidance in mosquito larvae

**DOI:** 10.1101/2025.04.02.646203

**Authors:** Melissa Kelley, Shubham Rathore, Karthikeyan Chandrasegaran, Cassandra Herbert, Chloe Elizabeth G. Sabile, Taylor Wood, Judd Joves, Andrew Palacios, Emily Susanto, Melissa Uhran, Arturo Ledezma Ramírez, Shyh-Chi Chen, Joshua Tompkin, Khwahish Singh, Muhmmad S. Khalid, Clement Vinauger, Elke Buschbeck, Patrick A. Limbach, Joshua B. Benoit

## Abstract

The gut microbiome is a rich source of nutrients that are critical to the development and biology of eukaryotes. Transfer RNAs (tRNAs) are essential components of protein synthesis, and some chemical modifications to tRNA rely on the availability of microbiome-derived nutrients. In eukaryotes, the micronutrient queuosine (Q) is salvaged from the microbiome or diet and then incorporated into eukaryotic tRNA to influence the speed and efficiency of protein synthesis. Here, we examine the role of microbiome-derived Q in mosquito larval development and behavior. When mosquito larvae are grown with a microbiome incapable of synthesizing Q, there is a significant impact on tyrosine levels and processes, which correlate with defects in behavior and cuticle formation. Due to defects in movement and behavioral responses, Q-deficient larvae demonstrate impaired predator evasion, leading to higher instances of capture by predaceous beetle larvae. The broad effects of Q-deficiency in mosquito larvae highlight the importance of microbiome-derived nutrients for eukaryotic physiology and behavior.

## Introduction

There are continuously emerging roles for the microbiome in eukaryotes, particularly as a key source of micronutrients and vitamins for the host organism (*1*, *2*). Mosquitoes are vectors for deadly human diseases (*3*) and growing pesticide resistance requires constant development of alternative population control strategies (*4*). Temperature and breeding sites have been linked to mosquito gut microbiome composition (*5*–*7*), suggesting a link between the environment and mosquito biology (*8*). There is mounting evidence that manipulating the microbiome is a valuable and transient method of population control (*9*), as the presence of specific microbial residents is crucial to mosquito development, fecundity, vectorial capacity, and survival (*8*, *10*–*13*).

Transfer RNA (tRNA) are essential for decoding codons of messenger RNA to add the appropriate amino acid to the peptide chain during protein synthesis. As such, tRNA links the genetic code to functional proteins in the cell, and tRNA are an unexplored target for mosquito population control strategies. Post-transcriptional modifications are essential for tRNA function as they directly impact the speed and efficiency of protein synthesis (*14*). The modification queuosine (Q) is located at the wobble position of tRNAs for four amino acids. As a wobble modification, Q influences the decoding of codons for these amino acids. The enzymatic machinery to synthesize Q is only found in bacteria, while lacking in eukaryotes. Therefore, eukaryotes must salvage Q from their gut microbiome or dietary queuine (q) (*15*–*19*). Our recent studies have suggested that Q may be involved in blood-feeding and reproduction of mosquitoes (*20*, *21*). In Drosophilids, Q modification is correlated with early developmental gene expression (*22*). However, it is unknown how a microbiome unable to produce Q would affect tRNA modification status and the biology of mosquitoes.

Here, we investigate the role of Q in mosquitoes by directly altering the larval microbiome to lack the ability to synthesize Q. We identify that Q is predominantly obtained through the gut bacteria in larvae and demonstrate that Q is a potent influencer of the larval transcriptome, especially in enzymes related to tyrosine metabolism and downstream processes. As such, we observe defective cuticular tanning and reduced dopamine levels in Q-deficient larvae, resulting in increased predation susceptibility. Ultimately, we demonstrate the importance of Q in mosquito larvae and provide the first evidence of an ecological connection between Q and predation in eukaryotes.

### 1. Acquisition of Q is dependent on the microbiome composition in mosquitoes

Mosquitoes obtain their microbiome during the early aquatic larval stage, and it is a critical source of specific micronutrients, such as B vitamins (*23*, *24*). To evaluate the influence of microbiome composition on tRNA modifications, mosquito eggs were sterilized, and larvae emerged with a microbiome limited to ΔqueA mutant *E. coli* or the control strain (K12). The queA gene is essential for the production of the tRNA modifications epoxyqueuosine (oQ) and, subsequently, queuosine (Q) in bacteria (*25*, *26*) (**Figure 1A**). Q modification is notoriously enigmatic as it is not synthesized by eukaryotes but salvaged via diet and gut bacteria for eukaryotic tRNA (*16*, *27*). Therefore, a lack of this gene would prevent the synthesis of Q in the microbiome and potentially affect the Q availability in mosquito tRNA.

**Figure 1.**
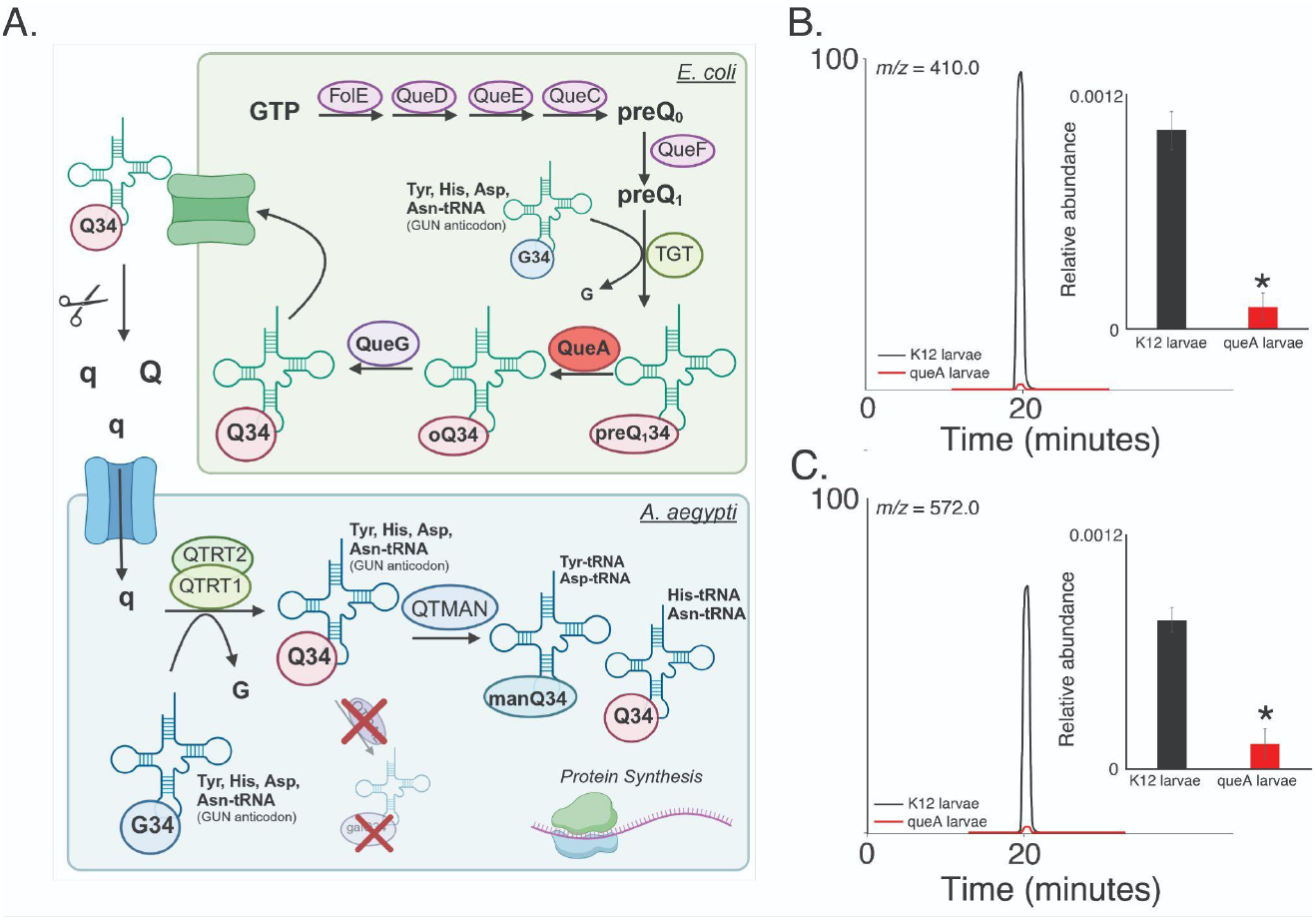
Queuosine (Q) is reduced when mosquitoes are reared with bacteria incapable of Q production. **A**. Mechanism for Q synthesis in bacteria and Q salvage in eukaryotes such as mosquitoes. **B**. Total ion chromatogram (TIC) of Q in tRNA of mosquito larvae with (K12, black) and without Q (ΔqueA, red) in the microbiome and the relative abundance of Q as determined by chromatographic peak area. **C**. TIC and relative abundance of the Q hypermodification, mannosyl-queuosine (manQ), in larvae reared with (K12, black) and without Q (ΔqueA, red) in the microbiome. Student’s t-test * P < 0.05. N = 3 with tRNA isolated from 20 larvae per replicate.

Expectedly, Q levels are significantly diminished in larvae grown with a ΔqueA microbiome (**Figure 1B**). In agreement with previous reports (*28*), the larvae have similar viability and development rates, regardless of Q levels (**Figure S1**). In eukaryotes, Q may be glycosylated to form mannosyl-queuosine (manQ) or galactosyl-queuosine (galQ) (*29*). As we previously established (*20*), only manQ is present in mosquito tRNA, and these levels were also lowered in the ΔqueA larvae (**Figure 1C**). Methylations have also been reported to be impacted by microbiome composition and Q availability in eukaryotes (*30*–*32*); however, only one modification, 2’-O-methyluridine (Um) was observed to be in higher abundance in ΔqueA larvae. The remaining modifications were at similar levels regardless of microbiome type (**Table S1**), which indicates that the larval tRNA epitranscriptome is relatively constant and largely independent of Q in the microbiome. Altogether, these results demonstrate that the bacterial composition of the aquatic larval environment directly influences Q modification in mosquito tRNA.

### 2. Dysfunction in gene transcription for ΔqueA larvae is recovered by q supplementation

There is a significant enrichment of transcripts related to cuticle formation, oxidoreductases, and fatty acid metabolism in ΔqueA larvae (**Figure 2A**). Downregulated transcripts in ΔqueA larvae were primarily involved in general metabolic processes, particularly in glycolytic and other sugar metabolic processes (**Figure S2**). To ensure Q levels were the underlying factor for transcriptional shifts, ΔqueA larvae were supplemented with dietary queuine (q), which resulted in a significant recovery of differentially expressed genes (**Figure 2B**). Approximately 80% of transcripts returned to control levels in the q-supplemented larvae (**Figure 2C**), demonstrating that Q is critical to the transcriptional profile of mosquito larvae.

**Figure 2.**
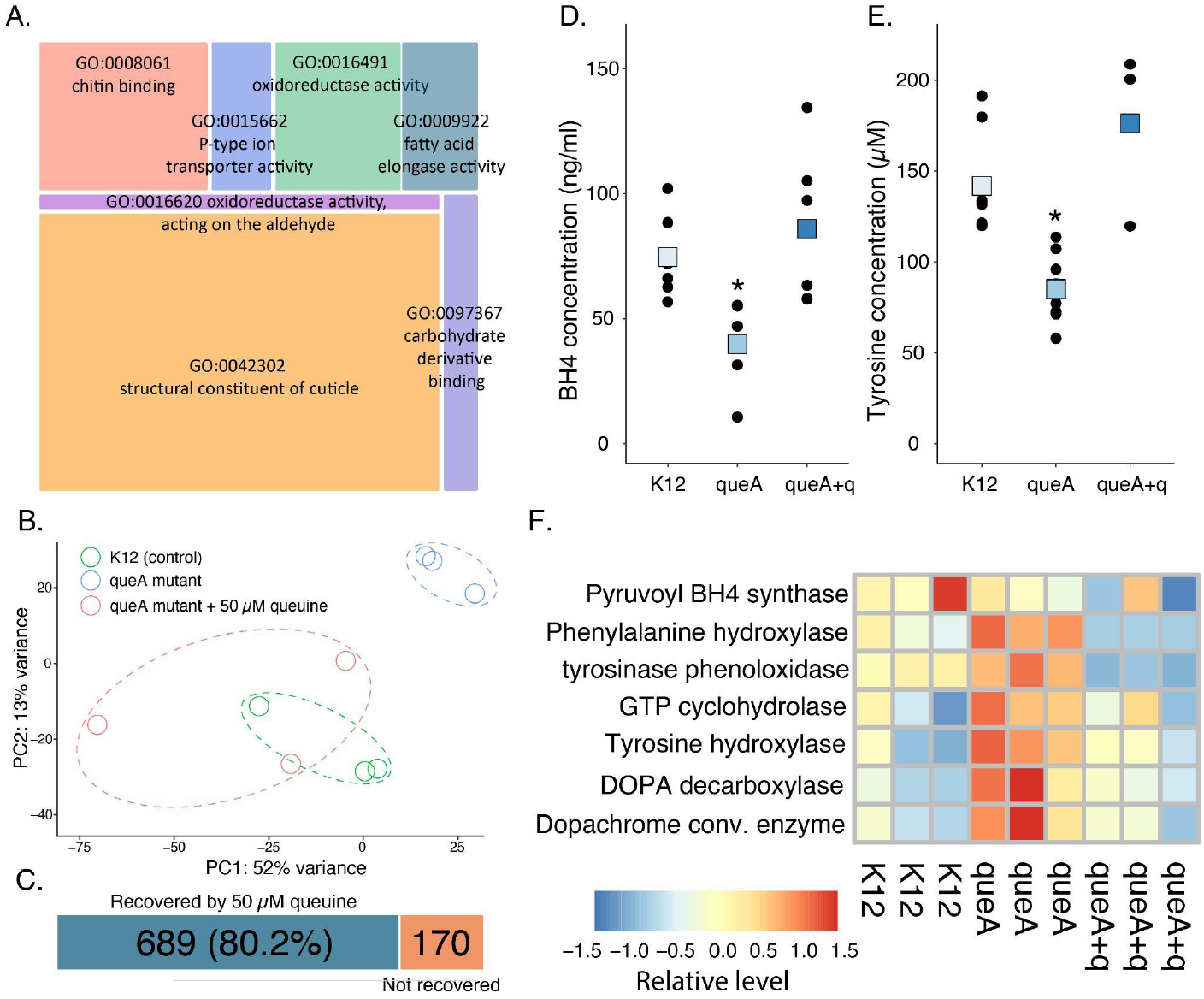
Transcriptional analysis indicated shifts in gene expression in tyrosine metabolism and downstream processes including cuticle development. **A**. Gene ontology analyses show that loss of microbiome-derived Q directly impacts the expression of cuticle and chitin-associated genes. **B**. PCA analyses of differentially expressed transcripts in K12, ΔqueA, and ΔqueA + q recovered larvae. **C**. General transcription patterns significantly shift following Q-deficiency and are recovered by q supplementation. N = 3 per treatment, FDR, P < 0.05. **D**. Larvae grown with less Q have lower tetrahydrobiopterin (BH4), which is required for the conversion of phenylalanine to tyrosine, general linear model, P < 0.05, N=9. **E**. Q-deficiency leads to a significant depletion of tyrosine, general linear model, P < 0.05, N = 9. **F**. Transcript levels for genes involved in the tyrosine-dopa-dopamine pathway are elevated in Q-deficiency.

Mice depleted of Q modification die from tyrosine withdrawal (*28*, *33*), raising the question of whether similar adverse effects are present in mosquito larvae. In ΔqueA larvae, tyrosine levels are lower and are recovered by q supplementation (**Figure 2D**), demonstrating a connection between Q and tyrosine levels which has been observed previously (*34*). In Q-deficient mice, tyrosine levels are lower due to the oxidation of the cofactor BH4, which is a cofactor for the enzyme that converts phenylalanine to tyrosine (*28*, *35*). Similarly, BH4 levels are reduced in Q-deficient mosquitoes (**Figure 2E**), indicating this is likely to contribute to lower tyrosine. Given that tyrosine is a precursor to many molecular processes (*36*, *37*), we investigated whether the microbiome composition impacted transcripts associated with the downstream pathways (**Figure S3**). Indeed, several enzymes involved in the tyrosine-dopa-dopamine pathway, which are critical in cuticle darkening and dopamine production, were enriched in ΔqueA larvae compared to control and recovered in ΔqueA larvae supplemented with q (**Figure 2F, Figure S4**). Ultimately, Q modification influences tyrosine levels and the enrichment of transcripts associated with these processes is likely compensatory for lower Q.

### 3. Q-deficient microbiome causes larval cuticle variation and behavioral differences

Considering the apparent transcriptional effects on tyrosine-related processes such as cuticle structure and darkening (*38*), we next evaluated how lower Q levels in the microbiome contribute to cuticle darkening. Larvae grown with ΔqueA bacteria consistently demonstrated lighter coloration throughout the organism (**Figure 3A**). Furthermore, the coloration differences were recovered when 10μM or 50μM q was supplemented to ΔqueA larvae (**Figure 3B**). Since q supplement recovers tyrosine levels as well, we assessed if a tyrosine supplement would also resolve the phenotype, which indeed was the case (**Figure S5**).

**Figure 3.**
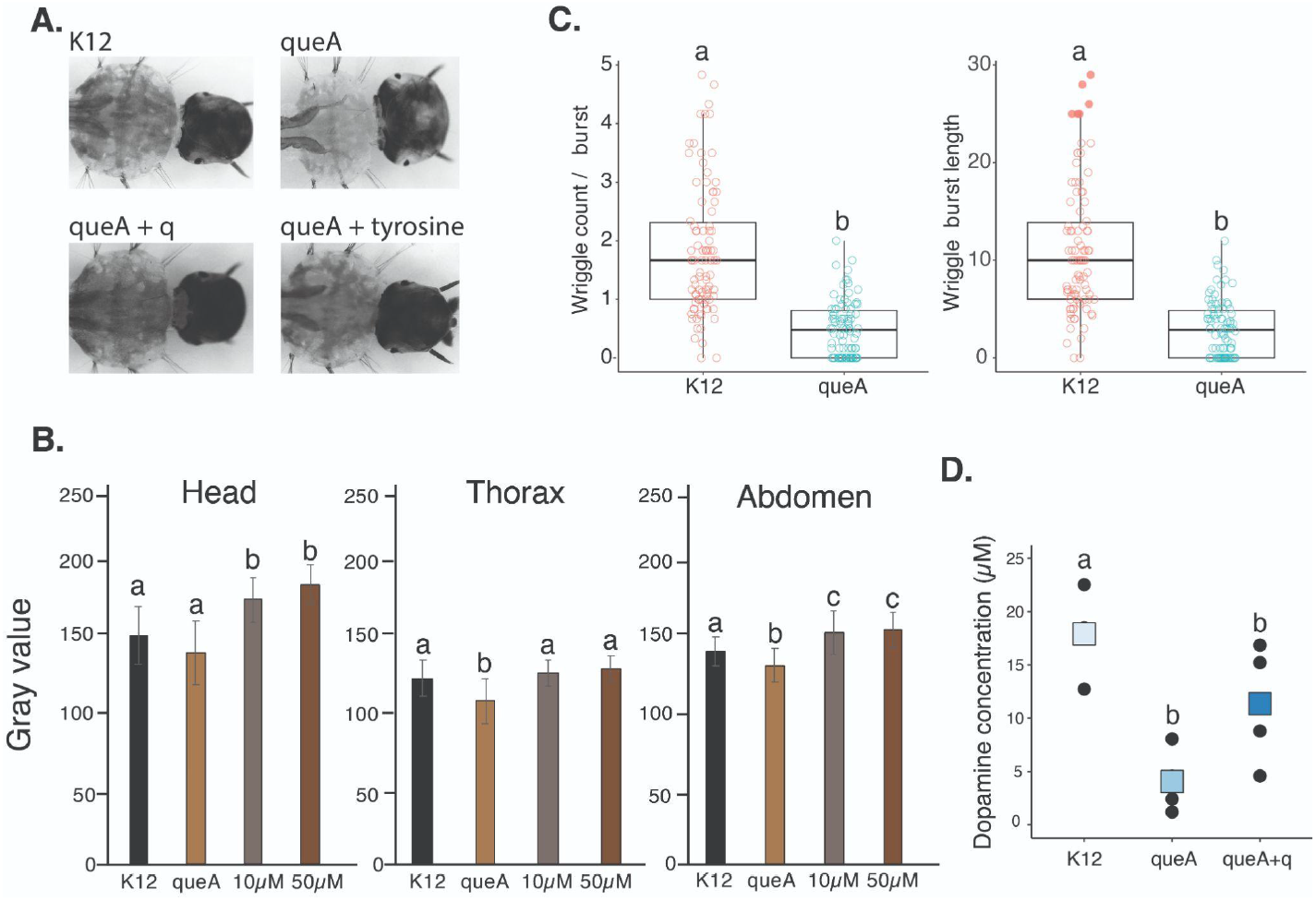
Cuticle darkening and behavior are affected by Q depletion in the microbiome of mosquito larvae. **A**. Representative images of cuticle darkening changes during Q-deficiency and recovery following supplementation with q or tyrosine. **B**. Quantification of darkening changes in the head, thorax, and abdomen was calculated by gray value. N = 15 samples per treatment. ANOVA, P < 0.05. **C**. Behavioral shifts in Q-deficient larvae based on wriggle count per burst and length of larval wriggle length, ANOVA, P < 0.05. **D**. Dopamine levels are lower in ΔqueA larvae and trending towards recovery in q supplemented larvae. General linear model, P < 0.05, N = 12.

The cuticle is essential for movement and structural integrity, and thus is critical for the wriggle burst motion characteristic of mosquito larvae (*39*, *40*). The Q-deficient larvae were found to have a smaller ratio of wriggles per burst and shorter wriggle burst lengths (**Figure 3C**). While the general wriggle burst counts and activities were similar (**Figure S6**), the shorter movements and fewer wriggles per burst imply behavioral differences in Q-deficient larvae. In Q-deficient mice, there are sex-dependent neurocognitive impairments, with female mice having issues in learning and memory formation (*41*). Dopamine is also a product of tyrosine and is implicated in arousal (*42*), movement (*43*), and environmental and stress responses in insects (*44*, *45*). Dopamine levels were lower in ΔqueA larvae (**Figure 3D**), indicating neurotransmitters are likely impacted by Q deficiency, possibly due to compromised tyrosine levels. Therefore, microbiome-derived Q is associated with lower tyrosine levels, which have downstream effects on cuticular darkening and locomotor activity in mosquito larvae.

### 4. Q in the microbiome is beneficial for predator evasion

Predator evasion is vital for the survival of mosquito larvae in natural habitats. With such effects in cuticular darkening, we next asked if the coloration would influence predation in environments with differing background contrasts. To test this, we subjected mosquito larvae to predation by a visually-guided ambush predator (*Thermonectus marmoratus*) with an extensively studied visual system (*46*, *47*). Beetle larvae were allowed to hunt mosquito larvae in assemblies with a white or black background (**Figure 4A**). Against a white background, the control (K12) larvae were predated significantly more, with only °50% of mosquito larvae surviving (**Figure 4B**). However, nearly 75% of K12 larvae survived and successfully evaded predation against a dark background (**Figure 4B**), due to blending in with the darker environment. In contrast, Q-deficient larvae were predated similarly regardless of the background color, indicating ΔqueA predation is not only being influenced by larval coloration. Altogether, there is a protective effect of Q on predation of mosquito larvae in darker environments.

**Figure 4.**
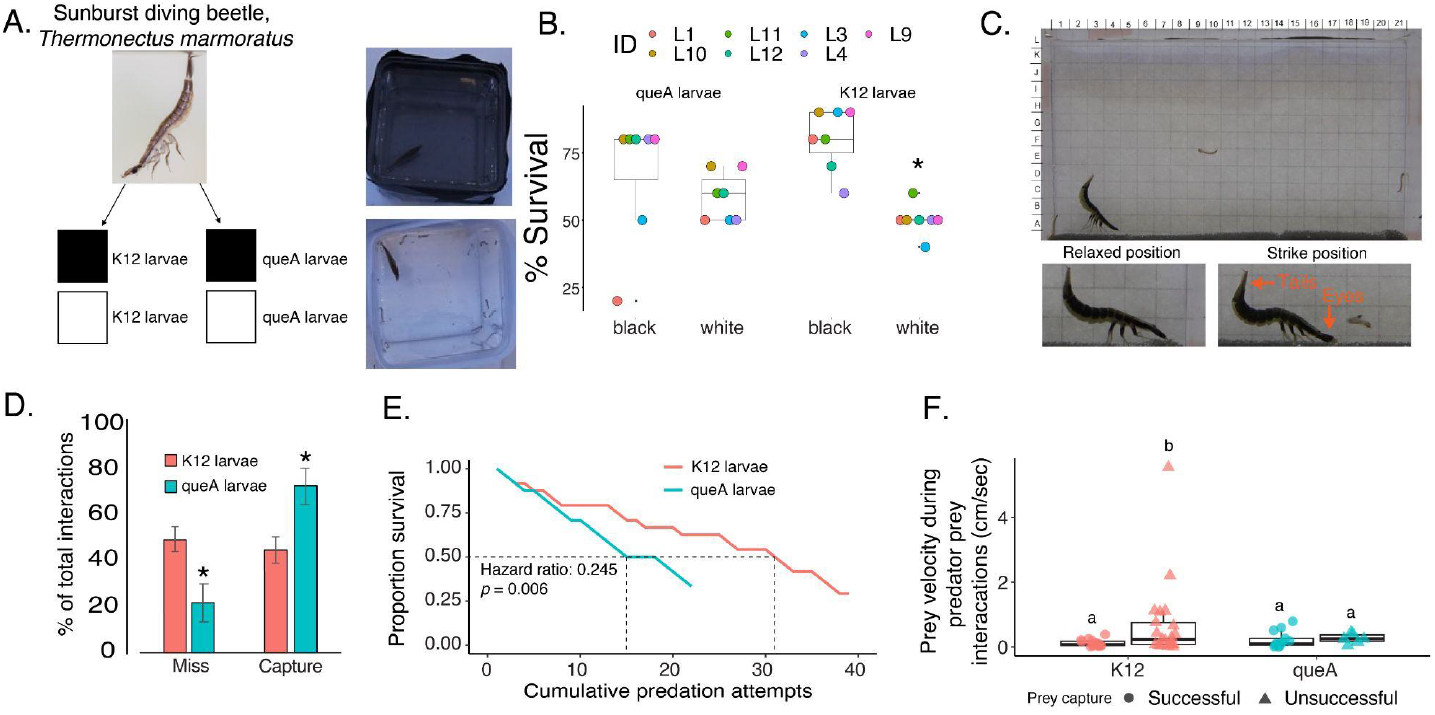
Predation is increased in Q-deficient mosquito larvae. **A**. Starved sunburst diving beetle larvae (*Thermonectus marmoratus*) were placed in arenas with white and black backgrounds and permitted to hunt mosquito larvae grown with ΔqueA and K12. Beetle larvae were given 30 minutes and 10 mosquito larvae were used in each assay. The same beetle larvae were used in each trial and given access to mosquito larvae and background in random combinations. **B**. Predation assays in these arenas show that K12 larvae are more susceptible to predation in the white background. ΔqueA larvae were predated regardless of background color. ANOVA, P < 0.05. N=7 with 10 mosquito larvae per assay. **C**. Predation assays were performed in a vertical arena with a grid background to allow the quantification of the hunting events. **D**. K12 and ΔqueA larvae have different miss and capture rates with beetle larvae capturing ΔqueA larvae with higher success. ANOVA, P < 0.05. N=8 with 3 mosquito larvae per assay. **E**. Compared to control mosquito larvae, ΔqueA larvae have lower survival with increasing predation attempts. N=8. ANOVA, P < 0.05, p = 0.006 **F**. K12 larvae that escape have a significantly higher escape velocity than ΔqueA larvae. General linear model, P < 0.05, N = 24 per treatment.

Since dopamine levels and movement are impacted by lower Q, we evaluated the relationship between predation and ΔqueA larvae further using a vertical predation assay (*46*). Our experimental assay allows for visualizing the ballistic strike of the predator to hunt the mosquito larvae (**Figure 4C**). The predator successfully captured Q-deficient larvae at a higher rate than control larvae (**Figure 4D**), and more predation attempts were necessary for the predator to capture control larvae (**Figure S7**).

Dopamine is involved in responding to environmental stressors and movement in insects (*43*), and lower dopamine in Q-deficient larvae may relate to an impaired capacity to evade a predator. Therefore, we asked if the Q-deficient larvae escaped predation with the same velocity as control larvae. When the predator strike was unsuccessful, the control mosquito larvae had a higher escape velocity, indicating an active attempt to evade the predator (**Figure 4F**). In contrast, ΔqueA larvae had similar velocity pre- and post-predation attempts regardless of capture outcome, suggesting Q-deficiency compromises mosquito larvae predator evasion. To ensure the effect was not caused by changes in the predator’s behavior, the time and distance to strike were found to be exact (**Figure S8**). When the predator was unsuccessful at capturing ΔqueA larvae, a spatial separation between the predator and prey influenced the interaction (**Figure S9**). This effect is not observed in control larvae, indicating that unsuccessful attempts to capture control larvae are driven by the prey escaping and supporting Q as protective in evasion from the predator. Taken together, the importance of Q from the microbiome includes significant molecular effects and extends to the ecological success of mosquito larvae in their environment.

### Summary

In this study, we demonstrate the tRNA modification queuosine (Q) is predominantly obtained from the microbiome, which is acquired from the aquatic environment of mosquito larvae. The availability of Q influences tyrosine levels through the cofactor BH4, which is critical in the conversion of phenylalanine to tyrosine. The reduction in tyrosine levels has downstream effects on dopamine biosynthesis and cuticular darkening. In Q-deficient larvae, there is compensatory expression of transcripts for enzymes involved in the tyrosine-dopa-dopamine pathway. Substantial recovery is observed when Q-deficient larvae are supplemented with dietary queuine (q), demonstrating the importance of this micronutrient on these processes. Above all, our findings indicate that Q is protective to mosquito larvae during predator evasion, implicating this micronutrient as a novel target for mosquito population control strategies during early development. Our results provide the first indication that Q improves mosquito fitness in predator-prey interactions through the modulation of components in the tyrosine-dopa-dopamine pathway.

Given the widespread evolutionary conservation of Q amongst eukaryotes and the diversity of bacteria in the gut microbiome (*15*), it may be possible that the microorganism composition of the larval aquatic environment causes fluctuations in Q availability in varying ecological circumstances. Microbiome surveys in mosquitoes show significant variation in the microorganism composition (*7*, *48*, *49*), and these bacteria have different capacities to utilize and synthesize Q (*50*). Taken together, Q levels are likely to vary among mosquito populations and larval environments. In our laboratory conditions, we observe increased predation of mosquito larvae with the loss of Q; however, follow-up studies will be necessary to evaluate defects in field environments. Thus, this study implicates the Q micronutrient as a complex influencer in the behavioral and ecological dynamics of insects and other animals.

## Supplemental Information

### Materials and Methods

### Mosquito rearing and *Escherichia coli* culturing

Mosquito eggs were collected from laboratory-reared *Aedes aegypti* mosquitoes. Rearing conditions have been described previously (*21*). *E. coli* cultures were obtained from *E. coli* Genetic Stock Center (Yale University (*51*)). The parental strain K12 (BW25113) was grown in Luria broth (LB) at 37°C, 170 rpm. The ΔqueA strain (JW0395) (*17*, *52*) was grown in LB medium with 30 ug/mL kanamycin at 37°C,170 rpm. Before larval culturing, the *E. coli* cultures were evaluated using OD600 and diluted to OD600 = 0.6 or approximately 4.8 x 10^8^ cells/mL using 1xPBS.

### Larval rearing with Q-deficient *E. coli*

The absence of Q modification was verified in the ΔqueA *E. coli* mutant tRNA using LC-MS/MS (**Figure S10**). Eggs are sterilized by submersion in 70% ethanol for five minutes, followed by 1% bleach for five minutes, and returned to 70% ethanol for five minutes (*23*). Immediately after, the eggs were placed into sterile tissue culture flasks (CellTreat info) with the appropriate *E. coli*. For all flasks, 3mL of diluted *E. coli* were added for a total of about 1.4 x 10^9^ cells or a concentration of 1.6 x 10^7^ cells/mL per flask. Approximately 500mg of autoclaved fish food (Tetramin) was added to each flask every two days or as needed. Mosquito larvae were visually assessed daily for survival and general growth. After 5-7 days post-emergence, larvae of the same instar (3rd/4th) were collected and washed with 1xPBS before use in subsequent experiments.

### Larval phenotype measurements

The larval phenotype was assessed approximately 5-7 days post-emergence or at the 3rd and 4th instar. Larvae were washed twice with 1xPBS and placed in a glass dissection dish with 1xPBS on ice for about 10 minutes to reduce movement before imaging (Dinolite, default settings). Larvae were imaged individually by placing the cold larva on a glass microscope slide and removing excess liquid. At least two images of each larva were taken to ensure the whole body was imaged. To process gray value, images were inverted and converted to grayscale using ImageJ to quantify the coloration. At least twenty points were placed randomly throughout the segment of interest and measured to generate a gray value; therefore, the larger the inverse gray value, the darker the coloration. Data was collected from four biological replicates with at least 15 larvae imaged for each treatment.

### Determining Q levels with liquid chromatography-tandem mass spectrometry

Prior to RNA isolation, approximately 20 mosquito larvae per biological replicate were washed with 1xPBS twice and frozen at −80 until further processing. Samples of tRNA were isolated and purified as previously described in adult mosquitoes (*20*, *21*) with 1.2μg being digested into nucleosides prior to LC-MS/MS analyses. Nucleosides were chromatographically separated as previously described (*20*), and detected using single reaction monitoring (SRM) on a Quantiva Triple Quadrupole mass spectrometer (Thermo Fisher Scientific). SRM transitions were used to identify the presence of a nucleoside (**Table S2**). To obtain a relative abundance, the TIC of transitions for specific modifications (i.e., 295 > 410 *m/z* for Q) were integrated and normalized with the summation of canonical nucleoside peak areas.

### Queuine supplementation and recovery of phenotype

Eggs were sterilized and inoculated as previously described. Two days post-emergence, mosquito larvae were placed in a sterile six-well plate (CellTreat) with 3-5 larvae per well. After washing with 1xPBS, ΔqueA larvae were placed in wells with varying concentrations of queuine (Toronto Research Chemicals, North York, Canada) at final concentrations of 10μM and 50μM. As controls, K12 and ΔqueA larvae were plated in tandem without q and treated with water as the vehicle. All larvae were monitored daily for survival and food availability. Approximately 100mg of autoclaved fish food was added to the wells every two days. At 5-7 days of exposure, larvae were removed from wells, washed with 1xPBS, and imaged as previously described.

Approximately nine larvae were imaged from each treatment. The experiments were repeated for a total of four biological replicates. Altogether, 25-30 larvae were imaged and quantified for coloration differences.

### RNA-sequencing of Q-deficiency in larvae

Total RNA was isolated using standard Trizol protocols and cleaned up using a GeneJet RNA kit. Library preparation and sequencing were performed by the Cincinnati Children’s Hospital and Medical Center DNA Core Facility. Libraries were prepared and sequenced using an Illumina NovaSeq 6000 system. The RNA-seq samples were processed according to our previous studies (*53*). Reads were mapped to predicted transcripts with Kallisto (*54*) under the default setting. Differential expression between contigs was examined with the DESeq2 package (*55*). A generalized linear model assuming a binomial distribution followed by the FDR approach was used to account for multiple tests (*56*) with Cut-off values for significance set at P < 0.05. Genes recovered following supplementation with q were determined based on expression reverting towards the control expression levels. After identifying the genes with differential expression, enriched functional groups were identified with gProfiler (*57*).

### Assays for tyrosine, BH4, and dopamine

Tyrosine levels were assessed using a tyrosine assay kit (Cell Biolabs). Three larvae per biological replicate were homogenized and subjected to the standard kit protocol. The assay was repeated with five biological replicates. Levels of BH4 were determined using a similar processing as in the tyrosine assay and quantified with a BH4 ELISA assay (Novus Biologicals). A dopamine ELISA kit (Abnova) was used to determine the dopamine level per larvae. All assays were conducted according to the manufacturer’s protocols with minor adjustments. Briefly, three larvae were homogenized, and the dopamine and BH4 was immediately extracted using the initial extraction protocol provided by the kit. Following this, the levels per larvae were assessed via competitive ELISA with four biological replicates (3 larvae each) for K12, ΔqueA, and ΔqueA + q treatment. Differences were assessed using generalized linear models (GLMs).

### Predator husbandry and vertical predation assay

*T. marmoratus* diving beetle larvae were collected from a lab-reared colony kept at 25°C on a 14h light - 10h dark cycle (*46*). One third-instar *T. marmoratus* was used per biological replicate, and nine total beetle larvae were evaluated in this assay. To test if the background affected predation, sunburst diving beetle larvae predate mosquito larvae at high rates (°10 larvae/hr) and were starved for 1 day before the assay. The assembly was covered with black or white paper, and each beetle larva was given 5 minutes to acclimate and 30 minutes to hunt 10 mosquito larvae. Beetle larvae were allowed to hunt larvae in assemblies with a white or black background (**Figure 4A**).

For the vertical assays, third-instar beetle larvae were starved for 24 hours prior to experiments to ensure hunting likelihood. The predation assays were performed in a vertical arena (10 cm × 1 cm × 8.5 cm) with a grid in the background to quantify in downstream video analyses. Each assay was 30 minutes long with 5 minutes of arena acclimation time. Each predator was given at least 30 minutes of rest between trials, and the same predator was subjected to both mosquito larvae types in a random sequence. Videos were manually assessed for predation time, predation attempts, and successes and further processed as described below.

### Analysis of larval movement and predator interactions

Video recordings (40 minutes each) were analyzed to study predator-prey interactions involving K12 and ΔqueA larvae (6 larvae with 1 predator per recording) based on previous methods (*39*, *40*). Every interaction was categorized as successful or unsuccessful, depending on whether the predator captured the prey. The behavior of both the predator and prey was quantified at three stages: immediately before, during, and immediately after the interaction. A predator-prey interaction was defined as the beginning when the predator set its eyes on the prey, which was identified by a distinct head bob and tail positioning change followed by directed movement toward the prey which marked the onset of the pre-capture phase, which lasted until prey capture. The prey handling time began at capture and ended when the prey was fully ingested. The total duration of a predation event was the sum of the pre-capture phase and the prey handling time. To assess prey behavior, wriggling activity was quantified through wriggle counts and wriggle burst length. The time larvae spent in various behavioral activities—browsing, filtering, resting, and wriggling—was also recorded. Additionally, prey velocity during interactions was measured to evaluate larval responses across successful and unsuccessful captures. Given the variability in predation event durations, all behavioral metrics were normalized. Predator approach distance was measured as the distance between the predator and prey when the predator noticed the prey. The time from prey detection to capture was recorded as the time to predator strike. The space-use patterns of predator and prey were tracked throughout each interaction to understand their spatial dynamics during the encounter. In parallel, larval behavior and spatial patterns were also quantified from video recordings where no predator was present, providing baseline activity levels for comparison. These baseline estimates allowed us to assess how larval behavior and space use were modulated in the presence of a predator. Generalized linear models (GLMs) were used to analyze predation attempts (Poisson regression) and success rates (binomial regression), with Tukey’s post-hoc tests for pairwise comparisons. Survival analysis evaluated larval survival over time using Kaplan-Meier curves and Cox proportional hazards models.

Behavioral responses were assessed using GLMs for wriggling activity and predator-prey distances. Differences in prey handling time and predator approach distances were analyzed using ANOVA, followed by Tukey’s tests for post-hoc comparisons. Linear mixed-effects models (LMMs) were used to analyze prey velocity and time to predator strike, accounting for trial-to-trial variability.

## Acknowledgments

These studies were supported by the National Institute of Allergy and Infectious Diseases of the National Institutes of Health under Award Number R01AI148551 and R21AI176098 (to J.B.B.), the National Institute of General Medical Sciences (R01 GM058843 to P.A.L.), National Institute of Food and Agriculture of the United States Department of Agriculture (Hatch project) VA-1017860 and VA-160212 (to C.V.), and National Institute of Allergy and Infectious Diseases of the National Institutes of Health under Award number R01AI155785 (to C.V.). Partial support was provided by the National Science Foundation IOS-1856241 (to E.K.B.).The content is solely the responsibility of the authors and does not necessarily represent the official views of the National Institutes of Health.

