## Supplemental materials for "A microbiome-derived nutrient underlies tyrosine metabolism and predator avoidance in mosquito larvae"

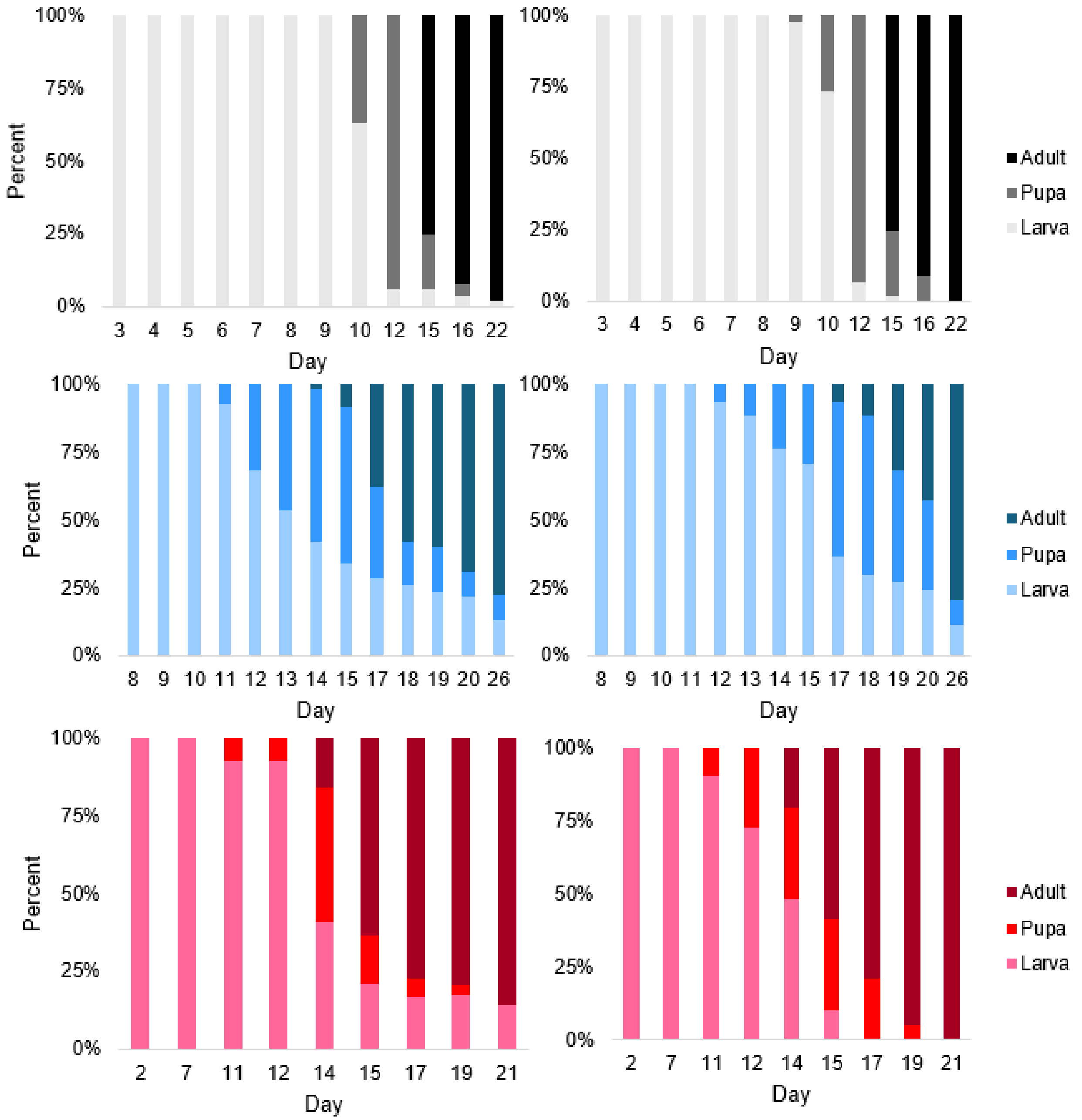


**Figure S1**. Transitions in developmental stage were similar in ΔqueA larvae and control K12 larvae. Three biological replicates demonstrate the time to pupation and adult emergence are aligned in the larva types.

**Table S1**. Relative abundance of modifications detected in mosquito larvae tRNA. The peak areas were normalized with the summation of the canonical nucleosides (C, U, G, A). A Student's t-test was used to determine significance. Only Q, manQ, and Um were significantly different between the larvae groups. Canonical nucleoside levels were the same across samples, demonstrating the amount of tRNA was the same.

| Modification | Q_L1 | Q_L2 | Q_L3 | K12_L1 | K12_L2 | K12_L3 | t-test |
|---|---|---|---|---|---|---|---|
| C | 0.11904 | 0.12491 | 0.12282 | 0.12166 | 0.12768 | 0.12054 | 0.730516 |
| U | 0.0318 | 0.0309 | 0.02989 | 0.03292 | 0.02939 | 0.03209 | 0.640027 |
| Ψ | 0.00464 | 0.00517 | 0.0051 | 0.00479 | 0.0056 | 0.00476 | 0.814576 |
| D | 0.00547 | 0.00652 | 0.00643 | 0.00579 | 0.0077 | 0.00575 | 0.721547 |
| $m^3C$ | 0.00581 | 0.01176 | 0.01632 | 0.00896 | 0.01056 | 0.01344 | 0.930352 |
| $m^5C$ | 0.02684 | 0.03054 | 0.03289 | 0.02917 | 0.03592 | 0.02888 | 0.6919 |
| Cm | 0.0102 | 0.01061 | 0.01035 | 0.01027 | 0.01161 | 0.01052 | 0.388944 |
| Um | 0.00057 | 0.00063 | 0.00054 | 0.00073 | 0.00071 | 0.00086 | 0.027236 |
| $m^5U$ | 0.00281 | 0.0032 | 0.00374 | 0.00312 | 0.00369 | 0.00309 | 0.895757 |
| Ψm | 7.9E-06 | 1E-05 | 1E-05 | 8.5E-06 | 1.1E-05 | 8.9E-06 | 0.986574 |
| A | 0.57925 | 0.5674 | 0.5521 | 0.56128 | 0.56787 | 0.56854 | 0.967868 |
| I | 0.00249 | 0.00276 | 0.00299 | 0.00252 | 0.00332 | 0.00255 | 0.878456 |
| Am | 0.00495 | 0.00322 | 0.00128 | 0.00393 | 0.00153 | 0.00399 | 0.997861 |
| $m^6A$ | 0.0102 | 0.01002 | 0.01212 | 0.01168 | 0.01239 | 0.00958 | 0.706777 |
| $m^1A$ | 0.11165 | 0.1304 | 0.13942 | 0.11859 | 0.15264 | 0.11943 | 0.836175 |
| $m^1I$ | 0.00253 | 0.00298 | 0.00361 | 0.00272 | 0.00341 | 0.00272 | 0.82051 |
| G | 0.26992 | 0.2768 | 0.29519 | 0.28414 | 0.27506 | 0.27882 | 0.879368 |
| $ac^4C$ | 0.00195 | 0.00246 | 0.00283 | 0.00224 | 0.00243 | 0.00253 | 0.953222 |
| $m^{6,6}A$ | 0.00446 | 0.0046 | 0.00437 | 0.00633 | 0.00436 | 0.00489 | 0.293099 |
| $m^2G$ | 0.01305 | 0.01501 | 0.01684 | 0.01321 | 0.01715 | 0.01353 | 0.850598 |
| Gm | 0.0073 | 0.00771 | 0.00821 | 0.00757 | 0.00786 | 0.00758 | 0.815447 |
| $m^1G$ | 0.03743 | 0.0422 | 0.04642 | 0.03781 | 0.04694 | 0.03864 | 0.831271 |
| $m^7G$ | 0.03912 | 0.04877 | 0.05702 | 0.04719 | 0.05497 | 0.04655 | 0.839003 |
| $ncm^5U$ | 1.8E-06 | 1.5E-06 | 1.8E-06 | 1.6E-06 | 1.8E-06 | 2.3E-06 | 0.443794 |
| $m^2_2G$ | 0.02031 | 0.02424 | 0.02909 | 0.02173 | 0.02492 | 0.02295 | 0.644551 |
| $mcm^5U$ | 0.00023 | 0.00025 | 0.00035 | 0.00025 | 0.00025 | 0.00029 | 0.652329 |
| $i^6A$ | 0.00282 | 0.00349 | 0.00446 | 0.00309 | 0.00296 | 0.00394 | 0.677593 |
| $acp^3U$ | 0.0011 | 0.00034 | 0.00023 | 0.00077 | 0.00037 | 0.00063 | 0.928232 |
| $acp^3D$ | 0.00013 | 2.7E-05 | 1.4E-05 | 8E-05 | 4.3E-05 | 4E-05 | 0.926351 |
| Q | 2.5E-05 | 3.8E-05 | 0.00031 | 0.00095 | 0.00103 | 0.00138 | 0.003623 |
| $t^6A$ | 2.8E-05 | 3.2E-05 | 4.5E-05 | 3.8E-05 | 4.9E-05 | 2.9E-05 | 0.668692 |
| oQ | 3.5E-07 | 4.3E-07 | 2.1E-05 | 1.1E-06 | 2.2E-06 | 1.6E-06 | 0.460256 |
| $m^6t^6A$ | 1.3E-06 | 1.6E-06 | 9.3E-07 | 7.3E-07 | 1.4E-06 | 1.3E-06 | 0.64714 |
| $ms^2t^6A$ | 5.2E-05 | 8.2E-05 | 7.2E-05 | 6.7E-05 | 3.8E-05 | 0.00016 | 0.642832 |
| manQ | 3.2E-05 | 5E-05 | 0.00037 | 0.00075 | 0.00084 | 0.00106 | 0.006841 |

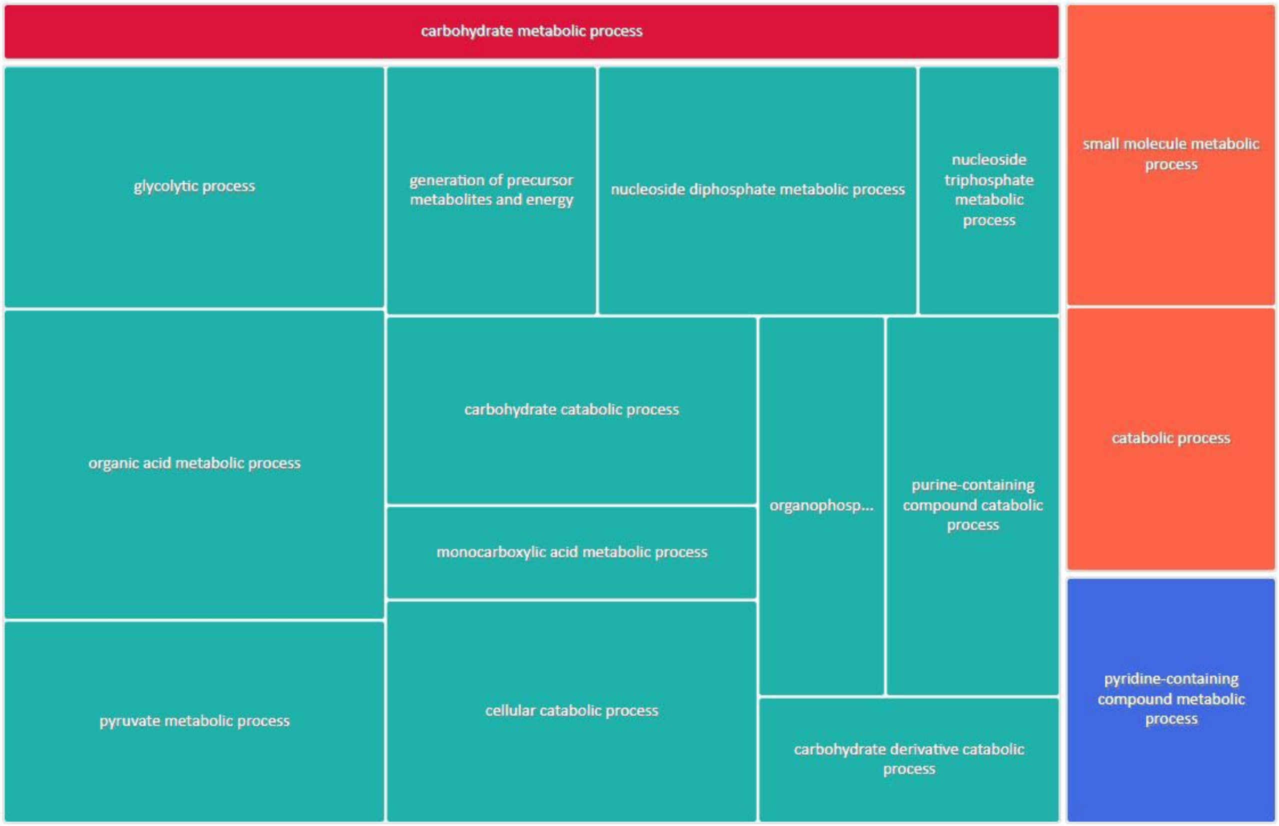


**Figure S2**. Transcripts that were downregulated in ΔqueA larvae compared to K12 controls such as catabolism, carbohydrate metabolism, and other metabolic processes.

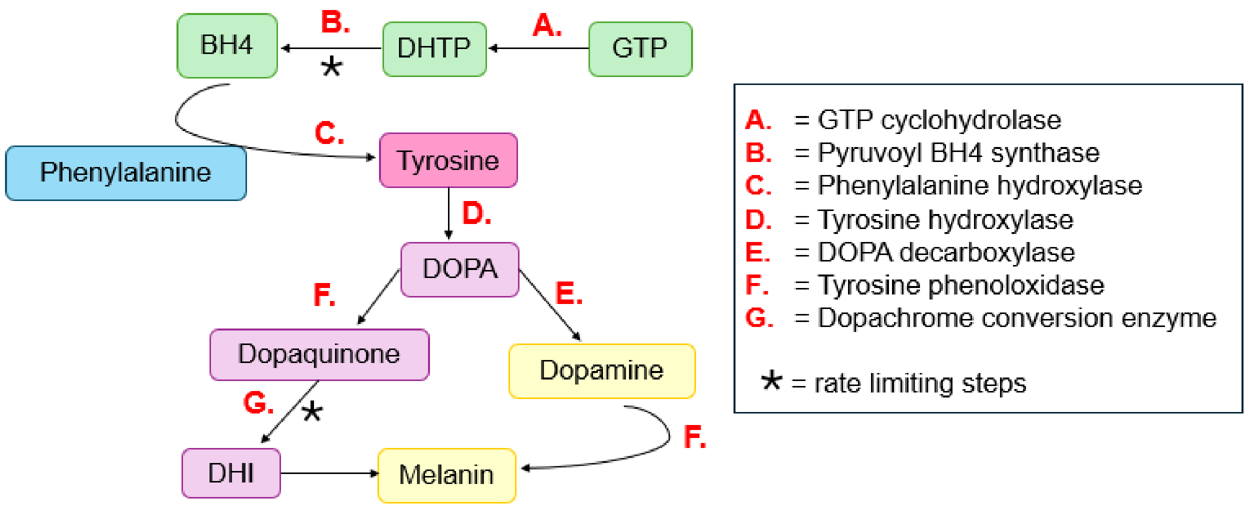


**Figure S3**. Pathways which rely on tyrosine as a precursor. Tyrosine is converted to L-dopa and then L-dopa can be converted to DHI or dopamine. The enzyme names and corresponding reactions are found here.

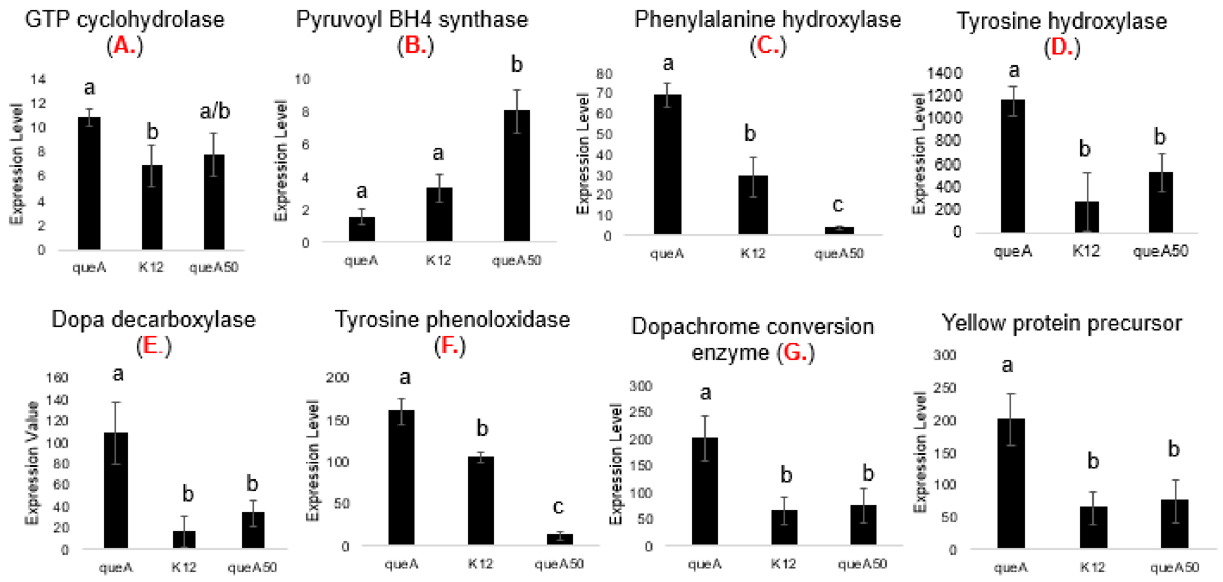


**Figure S4**. Expression values of specific enzymes involved in the tyrosine-dopa-dopamine pathway. Letters in red indicate pathway step shown in Figure S3. Expression values were determined significant with a p value < 0.05.

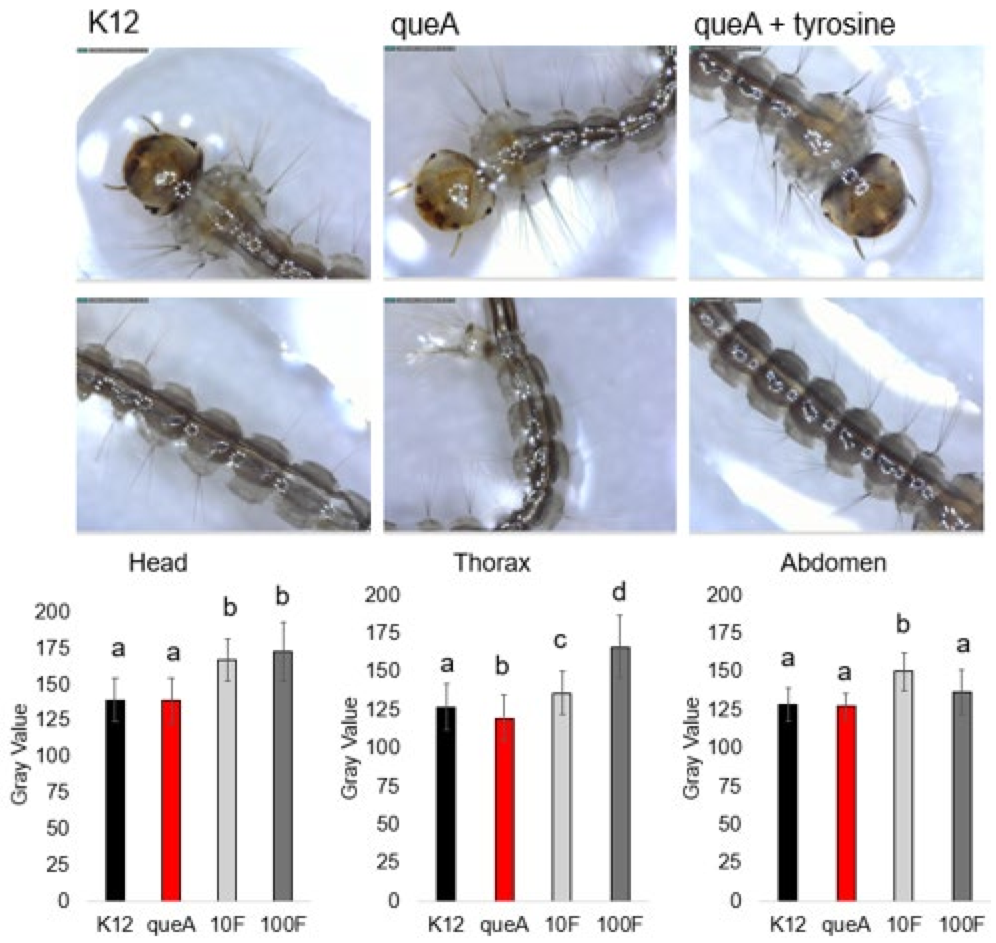


**Figure S5**. Tyrosine supplementation recovers the cuticular tanning defects observed in the ΔqueA larvae.

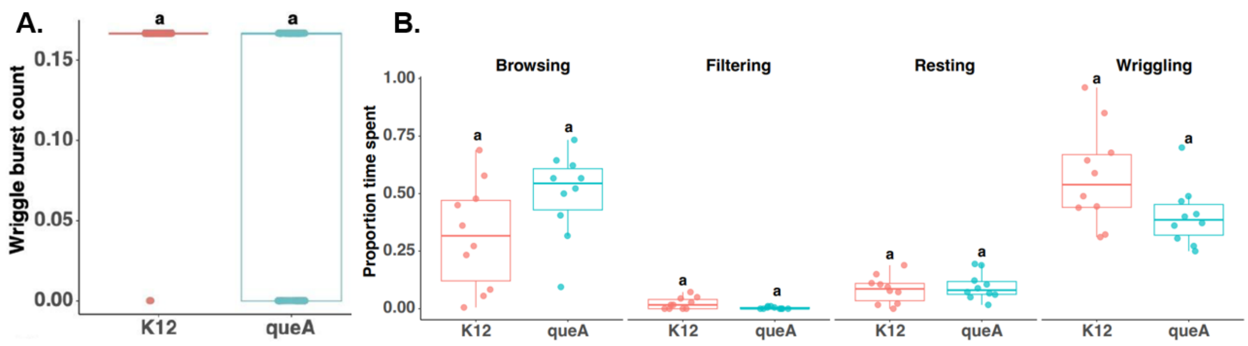


**Figure S6. A.** The overall wriggle burst counts were the same despite Q levels. **B.** The proportion of time spent on specific behaviors were the same regardless of Q levels in mosquito larvae.

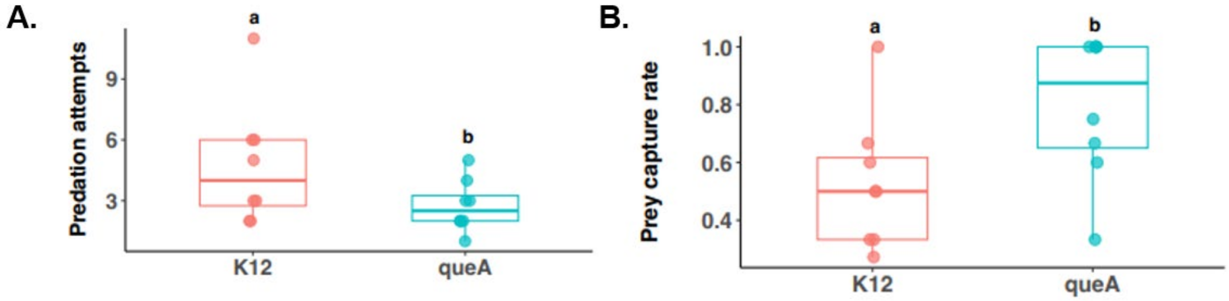


**Figure S7.** More predation attempts were necessary for the predator to capture K12 larvae. The prey capture rate was higher in ΔqueA larvae than K12, indicating queA larvae are captured more easily by the predator.

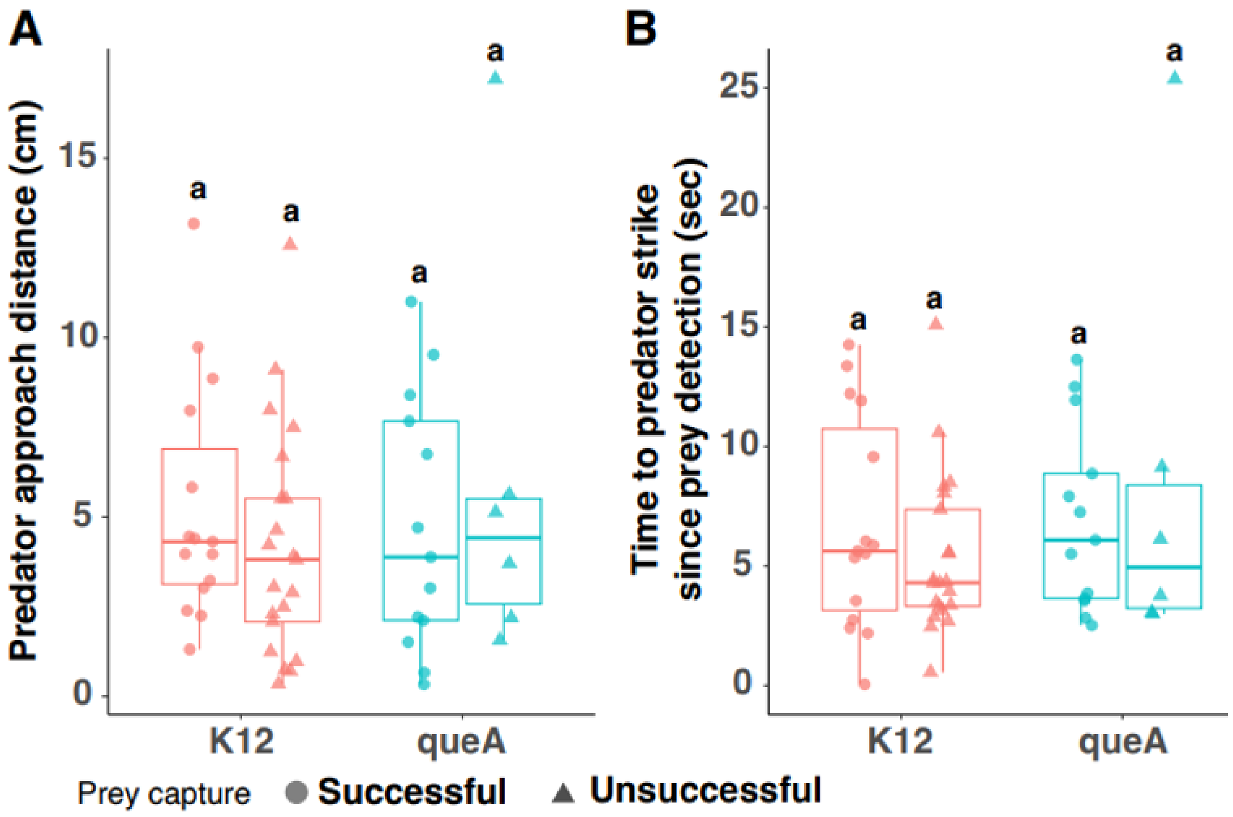


**Figure S8.** The predator approach distance and time to strike were similar regardless of Q levels in mosquito larvae, indicating the predator behavior is not altered during hunting.

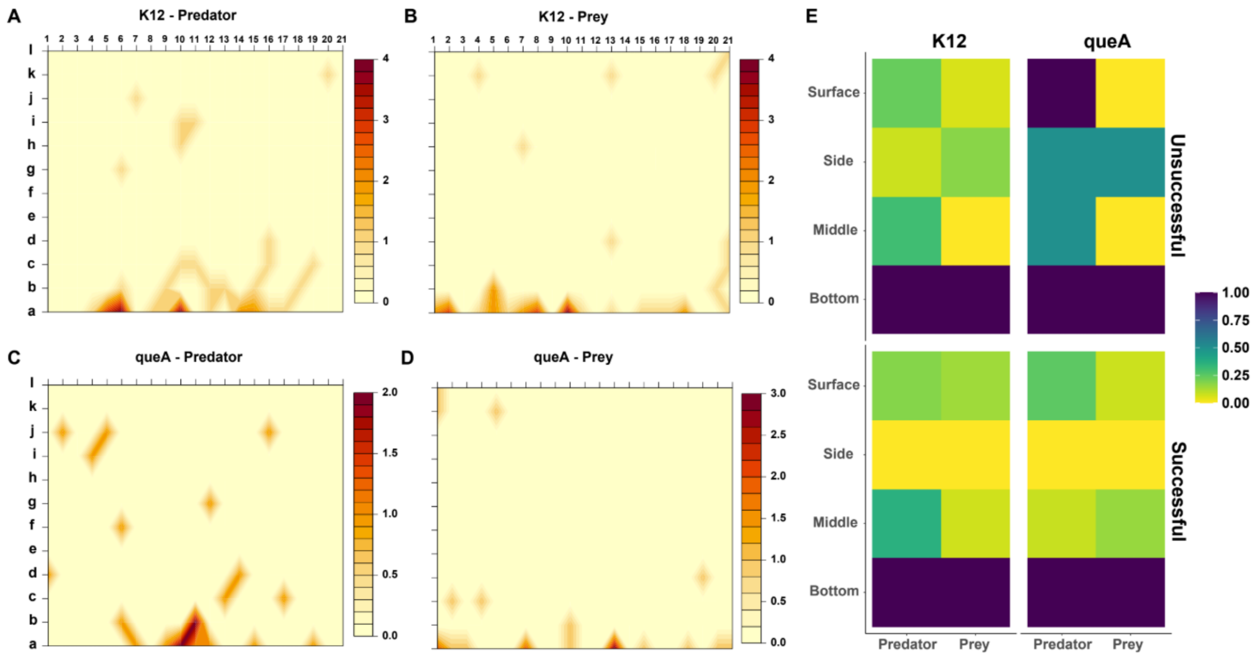


**Figure S9.** Predator-prey dynamics suggest that ΔqueA larvae were only found at the bottom of the vertical arena. This distinction in space use suggests that ΔqueA larvae that successfully evaded predators did so not through active escape behaviors but likely by taking advantage of non-overlapping spatial positioning with predators. ANOVA, $P < 0.05$. N = 24 per treatment.

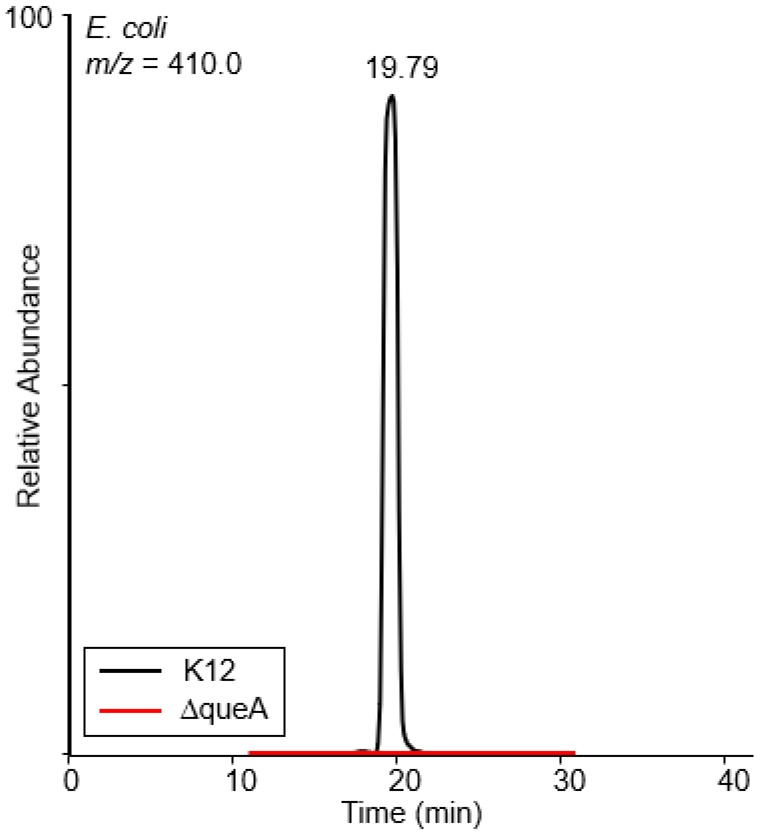


**Figure S10.** TIC of Q in the parental K12 *E. coli* strain (black) and the absence of Q in the ΔqueA strain(red) tRNA.

**Table S2**. Single reaction monitoring (SRM) transitions used for identification and quantification of nucleosides in mosquito larvae tRNA.

| Compound | Retention Time (min) | RT Window (min) | Polarity | Precursor (m/z) | Product (m/z) | Collision Energy (V) | RF Lens (V) |
|---|---|---|---|---|---|---|---|
| C | 2.48 | 3 | Positive | 244 | 112 | 12 | 41 |
| U | 4 | 8 | Positive | 245 | 113 | 10 | 30 |
| Y | 1.71 | 3 | Positive | 245 | 209 | 10 | 30 |
| D | 3 | 6 | Positive | 247 | 115 | 10 | 43 |
| Cm | 7.99 | 6 | Positive | 258 | 112 | 12 | 41 |
| m3C | 5.55 | 6 | Positive | 258 | 126 | 14 | 46 |
| m5C | 10 | 20 | Positive | 258 | 126 | 14 | 46 |
| m1Y/m3Y | 5.92 | 6 | Positive | 259 | 169 | 35 | 96 |
| m5U | 9.64 | 6 | Positive | 259 | 127 | 10 | 30 |
| Um | 14.17 | 6 | Positive | 259 | 113 | 10 | 30 |
| Ym | 5.72 | 10 | Positive | 259 | 223 | 10 | 30 |
| A | 23.3 | 10 | Positive | 268 | 136 | 17 | 63 |
| I | 9.23 | 6 | Positive | 269 | 137 | 10 | 40 |
| m5Um | 27.08 | 10 | Positive | 273 | 127 | 35 | 96 |
| m5Um | 27.08 | 10 | Positive | 273 | 241 | 35 | 96 |
| Am | 33.39 | 10 | Positive | 282 | 136 | 17 | 63 |
| m1A | 4 | 8 | Positive | 282 | 150 | 20 | 74 |
| m6A | 5.17 | 10 | Positive | 282 | 150 | 20 | 74 |
| m1I | 19.13 | 8 | Positive | 283 | 151 | 20 | 74 |
| G | 10.61 | 8 | Positive | 284 | 152 | 14 | 46 |
| ac4C | 22.68 | 6 | Positive | 286 | 154 | 10 | 46 |
| m6,6A | 33.67 | 8 | Positive | 296 | 164 | 28 | 79 |
| Gm | 20.43 | 10 | Positive | 298 | 152 | 11 | 50 |
| m1G | 23 | 20 | Positive | 298 | 116 | 16 | 50 |
| m2G | 30 | 33 | Positive | 298 | 166 | 16 | 50 |
| m7G | 8.65 | 10 | Positive | 298 | 166 | 16 | 50 |
| ncm5U | 5 | 8 | Positive | 302 | 170 | 35 | 96 |
| m2,2G | 31.2 | 10 | Positive | 312 | 180 | 15 | 60 |
| mcm5U | 23.39 | 15 | Positive | 317 | 125 | 10 | 30 |
| mcm5U | 23.39 | 15 | Positive | 317 | 153 | 10 | 30 |
| mcm5U | 23.39 | 15 | Positive | 317 | 185 | 10 | 30 |
| m2,2,7G | 31.21 | 10 | Positive | 326 | 194 | 31 | 96 |
| mcm5s2U | 34.06 | 20 | Positive | 333 | 141 | 35 | 96 |
| mcm5s2U | 34.06 | 20 | Positive | 333 | 169 | 35 | 96 |
| mcm5s2U | 34.06 | 20 | Positive | 333 | 201 | 35 | 96 |
| i6A | 36.94 | 6 | Positive | 336 | 136 | 31 | 96 |
| i6A | 36.94 | 6 | Positive | 336 | 148 | 31 | 96 |
| i6A | 36.94 | 6 | Positive | 336 | 204 | 31 | 96 |
| acp3U | 3.07 | 6 | Positive | 346 | 214 | 10 | 50 |
| k2C | 18 | 15 | Positive | 372 | 240 | 12 | 41 |
| ms2i6A | 41.41 | 8 | Positive | 382 | 182 | 31 | 96 |
| ms2i6A | 41.41 | 8 | Positive | 382 | 194 | 31 | 96 |
| ms2i6A | 41.41 | 8 | Positive | 382 | 250 | 31 | 96 |
| tm5U | 30 | 60 | Positive | 382 | 250 | 10 | 50 |
| ct6A | 34.81 | 6 | Positive | 395 | 263 | 31 | 96 |
| ms2io6A | 36.31 | 4 | Positive | 398 | 266 | 31 | 96 |
| tm5s2U | 36.74 | 15 | Positive | 398 | 266 | 35 | 96 |
| Q | 21.02 | 10 | Positive | 410 | 163 | 35 | 96 |
| Q | 21.02 | 10 | Positive | 410 | 295 | 35 | 96 |
| t6A | 31.43 | 10 | Positive | 413 | 281 | 31 | 96 |
| oQ | 16.5 | 7 | Positive | 426 | 163 | 29 | 84 |
| oQ | 16.5 | 7 | Positive | 426 | 295 | 12 | 84 |
| m6t6A | 32.49 | 4 | Positive | 427 | 295 | 31 | 96 |
| ms2t6A | 36.04 | 15 | Positive | 459 | 208 | 31 | 96 |
| ms2t6A | 36.04 | 15 | Positive | 459 | 327 | 31 | 96 |
| galQ | 25.85 | 10 | Positive | 572 | 163 | 35 | 96 |
| galQ | 25.85 | 10 | Positive | 572 | 295 | 35 | 96 |
| manQ | 22.94 | 10 | Positive | 572 | 163 | 35 | 96 |
| manQ | 22.94 | 10 | Positive | 572 | 295 | 35 | 96 |
| ISTD | 18.7 | 10 | Positive | 613 | 306 | 12 | 41 |
